# Inferring heterozygosity from ancient and low coverage genomes

**DOI:** 10.1101/046748

**Authors:** Athanasios Kousathanas, Christoph Leuenberger, Vivian Link, Christian Sell, Joachim Burger, Daniel Wegmann

## Abstract

While genetic diversity can be quantified accurately from high coverage sequencing, it is often desirable to obtain such estimates from low coverage data, either to save costs or because of low DNA quality as observed for ancient samples. Here we introduce a method to accurately infer heterozygosity probabilistically from very low coverage sequences of a single individual. The method relaxes the infinite sites assumption of previous methods, does not require a reference sequence and takes into account both variable sequencing errors and potential post-mortem damage. It is thus also applicable to non-model organisms and ancient genomes. Since error rates as reported by sequencing machines are generally distorted and require recalibration, we also introduce a method to infer accurately recalibration parameter in the presence of post-mortem damage. This method does also not require knowledge about the underlying genome sequence, but instead works from haploid data (e.g. from the X-chromosome from mammalian males) and integrates over the unknown genotypes. Using extensive simulations we show that a few Mb of haploid data is sufficient for accurate recalibration even at average coverages as low as 1-3x. At similar coverages, out method also produces very accurate estimates of heterozygosity down to 10^−4^ within windows of about 1Mb. We further illustrate the usefulness of our approach by inferring genome-wide patterns of diversity for several ancient human samples and found that 3,000-5,000 samples showed diversity patterns comparable to modern humans. In contrast, two European hunter-gatherer samples exhibited not only considerably lower levels of diversity than modern samples, but also highly distinct distributions of diversity along their genomes. Interestingly, these distributions were also very differently between the two samples, supporting earlier conclusions of a highly diverse and structured population in Europe prior to the arrival of farming.

---

The genetic diversity at a particular location in the genome is the result of its evolutionary past. Comparing the genetic diversity between individuals or regions of the genome thus gives insight into differences in their respective evolutionary histories. For a diploid individual, the heterozygosity of a genomic region (the fraction of sites in a region at which the individual carries two alleles) is the result of mutations that occurred since the two alleles shared a common ancestor. It is thus a function of the local mutation rate, but also genetic drift and selection, which affected the time that passed since the common ancestor. Variation in local mutation rates and, due to recombination, also in the strength of selection and genetic drift leads to variable diversity across the genome. Comparing heterozygosity between regions can thus identify locations that were affected differently by selection, or those with an increased mutation rate, while comparing heterozygosity between individuals may highlight differences in the demographic histories of populations.

While heterozygosity is readily obtained from high quality genotype calls by counting, it is much harder to infer accurately from low coverage genomes. This is primarily due to a substantial probability of observing only one of the two alleles and to sequencing errors, which occur at rates orders of magnitude higher than the expected heterozygosity in many species, including humans. Additional biases may be introduced by relying on a reference genome or by post-mortem DNA damage (PMD) when working with ancient DNA. A natural way of circumventing these issues is to infer genetic diversity probabilistically by taking many of the mentioned issues into account, and several such methods have been developed over the past decade. Johnson and Slatkin (2006), for instance, developed a method for estimating the population scaled mutation rate *θ* = 4*Nμ*, where *N* is the population size and *μ* the mutation rate, from large metagenomic data sets in the presence of sequencing errors. Shortly after, multiple moment based estimators were introduced to infer heterozygosity from a single individual (Hellmann *et al.* 2008; Jiang *et al.* 2008). Lynch (2008) then introduced a likelihood based estimator that relaxed the assumption of a known error rate by jointly estimating it together with heterozygosity from the data itself. Despite the additional parameter, this likelihood based estimator is generally more accurate, even if his implementation is ill-behaved at very low coverages (Lynch 2008).

Here we present a direct extension of the approach by Lynch (2008) that relaxes the assumption of infinitely many sites, does not require base frequencies to be known *a priori* and takes additional biases introduced by PMD fully into account. We achieve this by modeling genotype frequencies using the classic substitution model of Felsenstein (1981), which allows for back mutations, and by modeling PMD explicitly.

We further relax the assumption of a constant error rate. While variable error rates along and between individual reads is a well characterized feature of all current sequencing technologies, the provided estimates of these (the quality scores) are not reliable and must be recalibrated, particularly when coverage is low. This is commonly achieved by learning error rates from sites assumed *a priori* to be invariant, for instance by masking polymorphic sites, repetitive elements and large structural variants (DePristo *et al.* 2011). While we have extended this approach to tolerate PMD (Hofmanová *et al.* 2015), it requires detailed knowledge of the study species, which is often lacking for non-model organisms.

We circumvent this problem by using a reference-free recalibration approach that makes use of the base-quality information provided by sequencing machines. We rely on haploid sequences such as those from the X-Chromosome in male mammals and integrate over all possible but hidden genotypes while taking PMD and covariates such as position in read or read context into account. This renders our approach essentially free of reference biases since the reference is only required for aligning raw reads by mapping and current mapping techniques tolerate sequence divergence of up to 10% (e.g. Lunter and Goodson 2011).

Using computer simulations we show that our method reliably estimates local genetic diversity in single, diploid individuals even with average coverage below 2x for windows of ~ 1Mb. We further show that a few Mb of data at equally low coverage is sufficient to properly recalibrate distorted quality scores. Finally we use the here developed methods to infer the genome-wide pattern of diversity for several ancient and modern human samples. We found that these patterns differ between European and African samples, but that samples from a few thousand years ago cluster well with modern samples. In contrast, European hunter-gatherer individuals differ strongly from modern Europeans, but also from each other, illustrating the high diversity that existed in Europe before the neolithic transition.

## Theory

Here we develop a method to estimate heterozygosity from a collection of aligned reads by integrating out the uncertainty of the local genotype as well as the potential effects of postmortem DNA damage (PMD). Specifically, we are interested in inferring the stationary base frequencies ***π*** = {*π*_*A*_, *π*_*C*_, *π*_*G*_, *π*_*T*_}, along with the rate of substitutions *θ* = 2*Tμ* along the genealogy connecting the two alleles of an individual within a genomic region. Here, *T* corresponds to the time to the most recent common ancestor of the two lineages and *μ* to the mutation rate per base pair per generation. Notably, it is not possible to infer *T* and *μ* independently, and we therefore only attempt to estimate the compound substitution rate *θ* from the data.

To estimate *θ*, we will extend Felsenstein’s model of substitutions (Felsenstein 1981) to account for the uncertainty in the local genotypes. However, we will assume that base-specific rates of sequencing errors and PMD are known constants, motivated by the observation that rates of sequencing errors and PMD can be learned accurately from genome-wide data prior to inferring *θ* and ***π***, as we will show below.

### Inferring Heterozygosity

#### Substitution model

Let us denote the hidden genotype at site *i* by *g*_*i*_ where *g*_*i*_ consists of a pair of nucleotides *kl* with *k*, *l* = *A*, *G*, *C*, *T*. Under the substitution model, the probability of observing a specific genotype *g*_*i*_ = *kl* given the base frequencies ***π*** = {*π*_*A*_, *π*_*C*_, *π*_*G*_, *π*_*T*_} and the substitution rate *θ* is given by

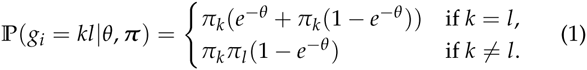

#### Emission probabilities

This model is easily extended to integrate out the uncertainty in observed genotypes. To do so we adopt a model similar to Lynch (2008) and those commonly used for genotype calling (Li 2011, e.g.). We will further closely follow the notation recently introduced by Hofmanová *et al.* (2015).

The observed data *d*_*i*_, at site *i* shall correspond to what is typically obtained when individual reads of next generation sequencing approaches were mapped to a reference genome. Here we will assume that all sequencing reads were accurately mapped and hence that reads with low mapping qualities have been filtered out. The data *d*_*i*_, obtained at site *i* thus consists of a list of *n*_*i*_ observed bases *d*_*i*_ = {*d*_*i*1_,…, *d*_*in*_*i*__}, *d*_*ij*_ = *A*, *C*, *G*, *T*.

We chose to model the observed data *d*_*i*_ at site *i* as a function of the underlying genotype *g*_*i*_ as well as the rates of sequencing errors and PMD, which we assume to be known for each observed base. Let us denote these base specific rates by *ϵ*_*ij*_ and *D*_*ij*_, respectively, for *j* = 1,…, *n*_*i*_ and further assume that the sequencing errors and PMD occur independently between reads. The likelihood of the full data at site *i* is thus given by

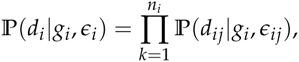

where *ϵ*_*i*_ = {*ϵ*_*i*,1_,…, *ϵ*_*i,n*_*i*__}.

Let us first develop the emission probability 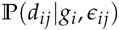 for the case of no PMD (*D*_*ij*_ = 0). Following Lynch (2008) and commonly used approaches (Li 2011, e.g.), we will assume that a sequencing read is equally likely to cover any of the two alleles of an individual and that sequencing errors may result in any of the alternative bases with equal probability *ϵ*_*ij*_/3. The probability of observing a base *d*_*ij*_ given the underlying genotype *g*_*i*_ = *kl* is then given by

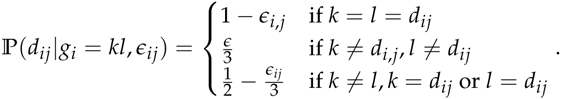

#### Post-mortem DNA damage

We will now extend this model with the possibility of PMD. The most common form of PMD is C deamination, which leads to a *C* → *T* transition on the affected strand and a *G* → *A* transition on the complimentary strand (e.g. Briggs and Stenzel 2007). These deaminations do not occur randomly along the whole read, but are observed much more frequently at the beginning of a read. This is due to fragment ends being more often single-stranded and thus subject to a much higher rate of deamination. Here we will develop our model for this form of PMD following the formulation of Skoglund *et al.* (2014), but we note that it is readily extended to incorporate other forms of PMD as well.

We feel that the rationale of our approach is best explained with a specific example. Consider *d*_*i,j*_ = *T* given the underlying genotype *g*_*i*_ = *CT*. There are three possible ways to obtain a *T*: i) by sequencing an allele *T* without error, ii) by sequencing an allele C affected by PMD without error, iii) or by sequencing an allele C not affected by PMD with error. We thus have

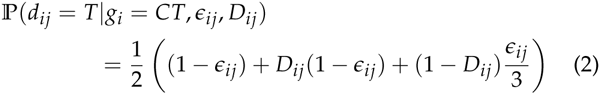

where *D*_*ij*_ denotes the probability that a *C* → *T* PMD occurred at the base of read *j* covering site *i*.

The emission probabilities for all combinations of *d*_*ij*_ and *g*_*i*_ derived following the same logic are found in the Appendix. Since we consider both *ϵ*_*ij*_ and *D*_*ij*_ to be known constants and in an effort to unburden the notation, we will refer to the emission probabilities simply as 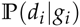 in the following.

#### Inference using EM-Algorithm

Assuming sites to be independent, the full likelihood of our model is given by

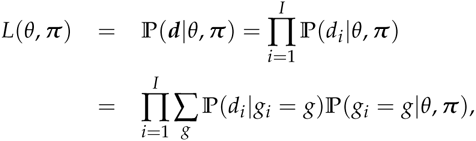

where the sum runs over all combinations *g* = *AA*, *AG*,…, *TT*.

To find the maximum likelihood estimate (MLE) of the model parameters *θ* and ***π***, we will adopt an Expectation-Maximization (EM) algorithm. The complete likelihood of our model is given by

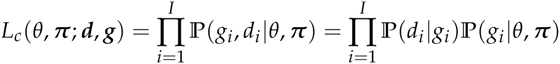

and thus the complete data log-likelihood by

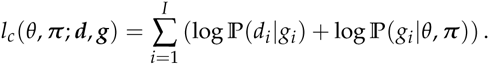

The expected complete data log-likelihood is calculated as

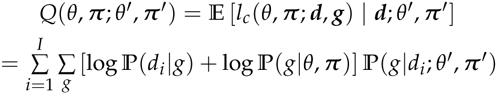

where the sum runs over all combinations *g* = *AA*, *AG*,…, *TT*. Only the second part *Q*_2_ of this sum depends on the parameters *θ*, ***π***. We have

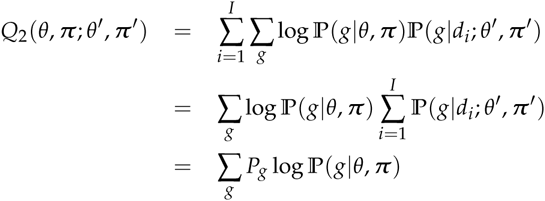

where we use the shorthand notation 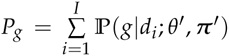.We have by Bayes’ Theorem

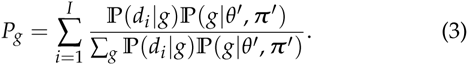

Let us write out *Q*_2_ explicitly:

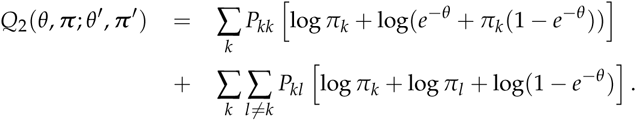

We have to maximize *Q*_2_ subject to the constraint

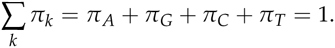

For this reason we form the Lagrangian

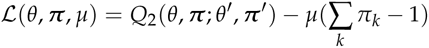

where *μ* is the Lagrange multiplier. We get the following partial derivatives of the Lagrangian:

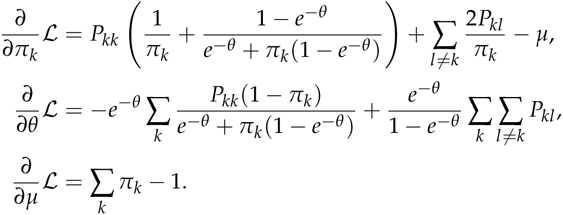

We have to set these equations to zero and solve for *π*_*k*_, *θ*, and *μ*.

With the parameter transformation *ρ* = *e*^-*θ*^ /(1 — *e*^-*π*^), the equations can be rewritten as (*k* = 1,…, 4):

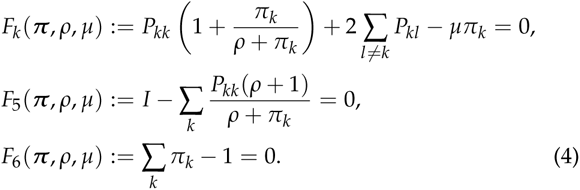

To streamline the notation, we will rename our variables:

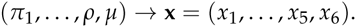

We will solve the above system

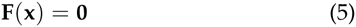

with the Newton-Ralphson method. We thus need to determine the 6 × 6 Jacobian matrix *J*_*ij*_ = ∂*F*_*i*_/∂*x*_*j*_. These are the non-zeros entries of the Jacobian where *k* = 1,…, 4:

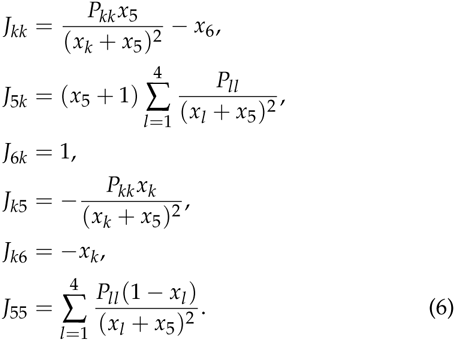

We can now approximate the zero of (5) with the iteration

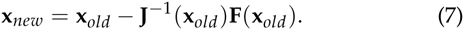

After a few iterations, we get the new estimate for the original parameters by setting *π*_*k*_ = *x*_*k*_ for *k* = 1,…, 4, and *θ* = −log(*x*_5_/(1 − *x*_5_)).

A brief outline of an efficient implementation of the algorithm is given in the Appendix.

#### Confidence intervals

We calculate an approximate confidence interval for *θ* using the Fisher information. To simplify the calculations we consider the *π*_*k*_ as constant. The observed Fisher information at the ML value 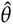 is

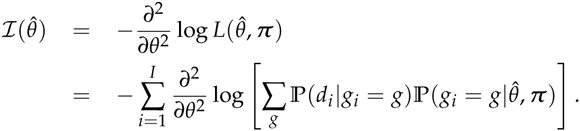

and the corresponding derivatives are

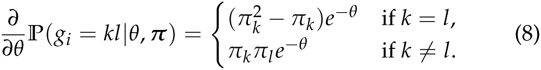

Observe that 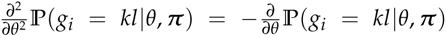. From this we easily get that

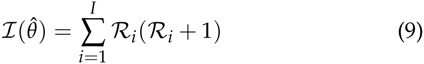

where we have set

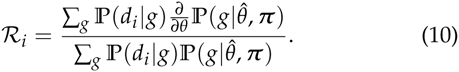

An approximate (1 — *α*) confidence interval is now given by

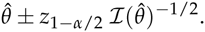

### Estimating base-specific error rates

The challenge of inferring genetic diversity from next-generation sequencing data lies in the fact that the per base error rates are orders of magnitude higher than the expected heterozygosity of many species (Lynch 2008). While this issue can easily be overcome with high coverages, accurate inference from low-coverage data relies on an exact knowledge of base-specific error rates. Crude estimates of these rates are usually directly provided by the sequencing machines themselves. However, these estimates are often inaccurate and are recommend to be recalibrated for genotype calling (DePristo *et al.* 2011).

The most commonly used approach for recalibration is BQSR (Base Quality Score Recalibration) implemented in GATK DePristo *et al.* (2011); McKenna *et al.* (2010). This approach infers new quality scores by binning the data into groups based on covariates such as the raw quality score, the position in the read or the sequence context. All bases within such a bin are assumed to share the same error rate, which can be readily inferred if the true underlying sequence is known. As an alternative, Cabanski *et al.* (2012) proposed to fit a logistic regression to the full data where the response variable is the probability of a sequencing error and the explanatory variables are the raw quality scores and covariates such as position in the read or base context.

For our purpose, these methods suffer from two shortcoming: first, they can not be applied to ancient DNA since they do not take PMD into account. Second, both require a reference sequence as well as knowledge on polymorphic positions such that they can be excluded from the analysis. While we have shown how to extend the BQSR method to ancient DNA (Hofmanová *et al.* 2015), we here develop an approach that also integrates over the unknown reference sequence.

To do so, we will assume that there exists a genomic region for which the individual does not show any polymorphism. A good example of such a genomic region are non-homologous sequences from sex chromosomes in heterogametic individuals (e.g. most of the X chromosomes in mammalian males), and we will describe our approach having this type of data in mind. However, we note that our approach is also readily applied to diploid regions that are known to be monomorphic, such as positions that are highly conserved among species or positions retained after filtering out those with high minor allele counts (Cabanski *et al.* 2012).

#### Model

As above, let us denote hidden genotype at site *i* by *g*_*i*_ where *g*_*i*_ is one of the nucleotides *A*, *G*, *C*, *T*. At each site *i* there are *n*_*i*_; reads and we denote by *d*_*ij*_, *j* = 1,…, *n*_*i*_ the base of read *j* overing site *i*. A sequencing error occurs with probability *ϵ*_*ij*_. These probabilities shall now be given by a model

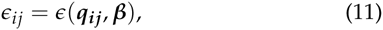

where ***q*_*ij*_** = (*q*_*ij*1_,…, *q*_*ijL*_) is a given external vector of informations and ***β*** = (*β*_0_,…, *β*_*L*_) are the parameters of the model that have to be estimated. While our approach is flexible regarding the choice of included covariates, we will here consider the raw quality score, the position within the read, the squares of these to account for a non-linear relationships, and all two-base contexts consisting of the bases of the read at positions *i* – 1 and *i*.

Following Cabanski *et al.* (2012) we impose the logit model

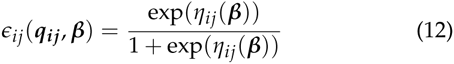

with

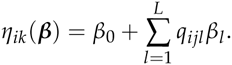

In the case of monomorphic or haploid sites only, the probability of the read vector *d*_*i*_ given the hidden state *g*_*i*_ can be written more generally as

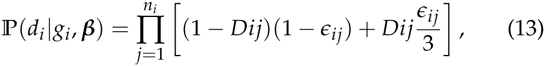

Here, the dependence on the parameters ***β*** is given by (12),

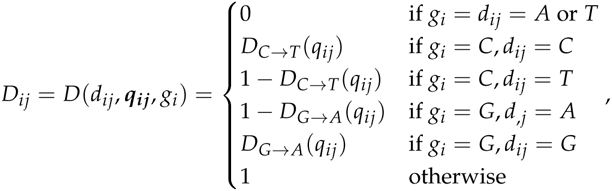

and *D*_*C* → *T*_(*q*_*ij*_) and *D*_*G* → *A*_(*q*_*ij*_) refer to the known probability that a *C* → *T* or *G* → *A* PMD occurred at the position covering site *i* in read *j*.

#### >EM-Algorithm

We propose an EM algorithms for this estimation that is similar to the one above, but assume here that the base frequencies *π*_*g*_, *g* = *A,G,C,T* are known, i.e. can be derived accurately from counting in the region. The complete data log-likelihood of our model is given by

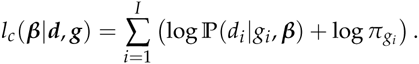

From this we get the expected complete data log-likelihood

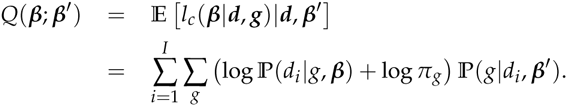

For the M-step we need only to consider the first part of *Q*(***β***; ***β***′):

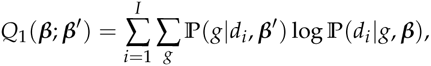

where

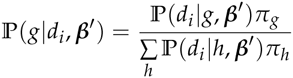

by Bayes’ formula. From (13) we get more explicitly

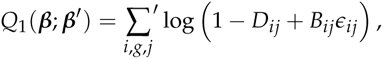

where we used the abbreviations 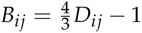 and

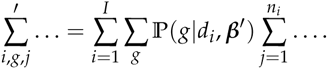

In order to maximize *Q*_1_ for ***β***, we calculate the gradient vector **F**(***β***) = ∇_***β***_*Q*_1_(***β***;***β***′) with components

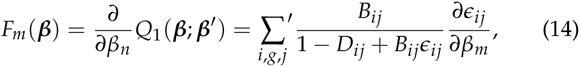

for *m* = 0,…, *L*. From (12) we obtain

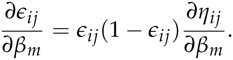

Observe that ∂*η*_*ij*_ /∂*η*_0_ = 1 and ∂*η*_*ij*_ /∂*β*_*m*_ = *q*_*ijm*_ for *m* = 1,…, *L*.

We solve **F**(***β***) = (**0**) with the Newton-Ralphson method with the Jacobian matrix *J*_*mn*_ = ∂*F*_*m*_ /∂*β*_*n*_. From (14) we get

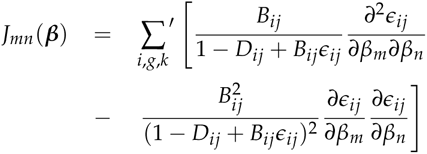

where

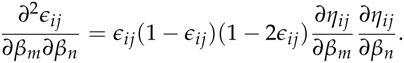

Putting everything together we obtain

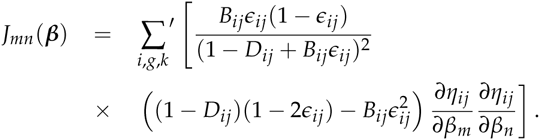

The Newton-Ralphson iteration is

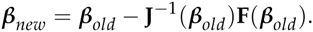

### Estimating Rates of Post-Mortem Damage

As mentioned above, the most common form of PMD is C deamination, which leads to a *C* → *T* transition on the affected strand, and a *G* → *A* transition on the complimentary strand (e.g. Briggs and Stenzel 2007). These deaminations occur more frequently in single stranded DNA, and are therefore observed more frequently close to natural break-points, i.e. at the ends of the DNA fragments. Consequently, the rates of PMD, while specific to the sample and the sequencing protocol used, are generally decaying roughly exponentially with distance from the ends of the read Skoglund *et al.* (2014). Since ancient DNA is highly fragmented, one read can often cover an entire DNA molecule, and hence *C* → *T* and *G* → *A* transitions may be seen in a single read, but are accumulated and opposite ends.

Here we follow Jónsson *et al.* (2013) and estimate PMD rates directly from genome-wide counts of *C* → *T* and *G* → *A* transitions as a function of distance within the read. For this we first build the three-dimensional table *T* where each entry *T*_*rsp*_ corresponds to the number of observed bases *r* read at a site with reference base *s* at position *p* within a read. While these counts depend on the divergence between the sequenced individual and the reference genome used for mapping, we here develop an approach that takes this divergence into account.

#### Position specific estimator

Let us denote by *μ*_*rs*_ the probability of a true difference between the sequenced individual and the reference such that the reference has base *r* and the sequenced individual base *s*. Since the reference and a sequenced chromosome form a genealogy on which these mutations occurred, it is safe to assume that *μ*_*rs*_ = *μ*_*sr*_. We will further assume that the observed counts in a cell *T*_*rsp*_ not affected by PMD are a direct function of *μ*_*rs*_. We thus have

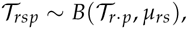

where B(·, ·) is the binomial distribution and

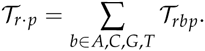

For cells affected by PMD, such as *T*_*CTp*_, we then have

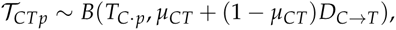

where *D*_*C* → *T*_ is the rate of *C* → *T* PMD.

Under the assumption that *μ*_*rs*_ = *μ*_*sr*_, we obtain an ML estimate of *D*_*C* → *T*_ (and analogously for *D*_*G*→*A*_) as

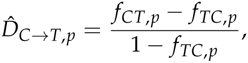

where

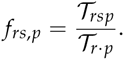

We note that this approach may lead to an ML estimate of D < 0 when *ƒ*_*CT*_ < *ƒ*_*TC*_. In this case we set D = 0, and our estimator thus corresponds to a maximum *a posteriori* estimate using a uniform prior 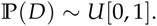.

We further note that this approach assumes that all differences observed between reads and the reference are due to divergence or PMD, but not sequencing errors. While sequencing errors, divergence and PMD can not be jointly inferred, additional insight into the accuracy of our approach is gained by studying the alternative extreme case in which all difference observed between reads and the reference are assumed to be due to PMD or sequencing errors alone. By denoting the genome-wide sequencing error rate by *ϵ*, the relevant equations become

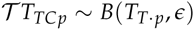

and

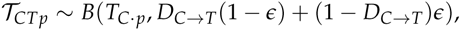

and the ML estimate

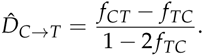

Since average sequencing error rates are on the order of 1%, they dominate the table *T* only in cases when *ƒ*_*TC*_ are small (on the order of 1%). As a consequence, the error when estimating the rates of PMD due to the omission of the factor of 2 in the denominator is never larger than 1% of the estimated value.

#### Exponential model

Since the rate of PMDs is generally low far away from the read ends, position specific estimates may become noisy for these positions, particularly if data is limited. We thus also introduce a method to estimate parameters of a model of exponential decay with the position in the read. The use of such a model was first introduced by Skoglund *et al.* (2014), and we implement here a slightly more general version of their function. Specifically, we will assume that the probability of observing base *T* when the reference sequence is a *C* at position *p* is given by

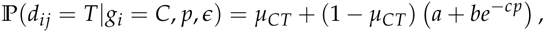

where *μ*_CT_ again denotes true differences between the individual and the reference.

To obtain ML estimates for the parameters of this probability function we again turn to the Newton-Raphson algorithm as shown in the following. However, we note that some of the parameters are non-identifiable, and we thus show here how to obtain estimates for the parameters of the probability function

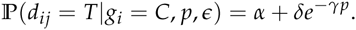

The log likelihood of the data is then given by

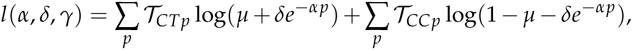

the gradient vector **F** (*α*, *δ*, *γ*) by

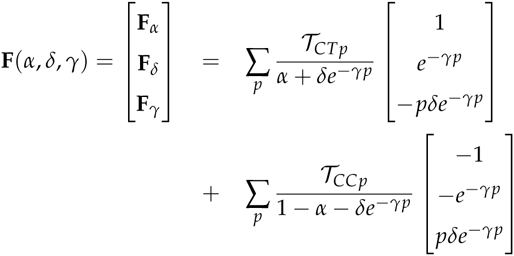

and the Jacobian matrix **J** (*α*, *δ*, *γ*) by

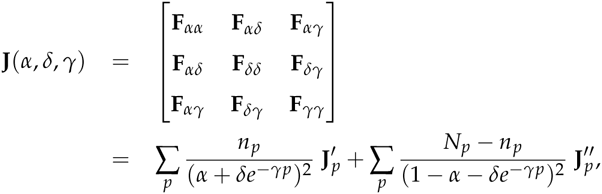

where

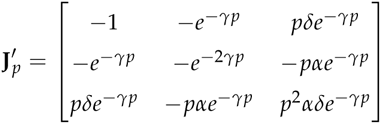

and

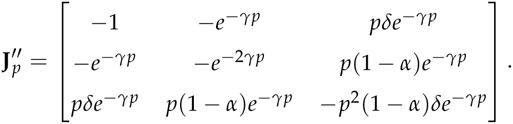

The Newton-Raphson iteration for ***θ*** = (*α*, *δ*, *γ*)^*T*^ is given by

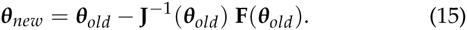

From these estimates we now obtain estimates tor our parameters *μ*_*CT*_, *a*, *b* and *c* as follows. First, and under the assumption that *α*_*CT*_ = *α*_*TC*_, we obtain the ML estimate

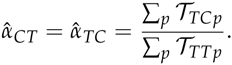

Then, 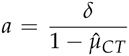, *b* = *γ* and 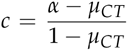.We use the analogous logic to infer PMD patterns for *G* → *A* damages, but measuring positions from the opposite endof the read.

### Implementation

All approaches mentioned were implemented in a custom C++ program available at our lab website. We used functions included in the library BamTools for manipulating bam files (Barnett *et al.* 2011).

## Simulations

### Generating simulations

In this section we illustrate the power and accuracy of our inference approaches with simulations. These were generated using a custom made R script that implements the following steps:

1. The first chromosome of length *L* was simulated using random bases with frequencies ***π*** = {0.25,0.25,0.25,0.25}.
2. The second, homologous chromosome was simulated according to the Felsenstein (1981) substitution model (eq. 1) with ***π*** and a chosen *θ* value.
3. Sequencing reads of 100 bases were then generated by copying from one of the two chromosomes with equal probability and by choosing a starting position uniformly between positions 1 – *L* and *L* until the desired average coverage was reached. All reads copied from the second chromosome were considered to map to the reverse strand.
4. Post-mortem damage (PMD) was simulated on all reads with probabilities following an exponential decay with increasing position in the read as proposed by Skoglund *et al.* (2014) to match realistic patterns. Specifically, we simulate PMD at position *p* within the read with probability

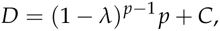

where *λ* = 0.3 and *C* = 0.01 for both *C* → *T* and *G* → *A* but with *p* counted from the 3′ and 5′ ends, respectively.
5. For each simulated base, a phred-scaled quality score was simulated and sequencing errors were then added with probabilities given by these scores. If not stated otherwise, quality scores were simulated from a normal distribution with mean *μ*_*Q*_ and standard deviation *σ*_*Q*_, truncated at zero. When testing our recalibration approach, however, the quality scores were simulated from a uniform distribution *U*[5,60] and then transformed according to eq. 12 with coefficients ***β*** to obtain the true error rate, with which sequencing errors were simulated.
6. The simulated data was finally used to generate a reference FASTA file containing the first chromosome and a SAM file containing the reads. The latter was then transformed into a BAM file using samtools Li *et al.* (2009).

### Power to infer Heterozygosity

To check the power of our approach to infer *θ* from low coverage data, we first simulated data within a 1 Mb window with a true *θ* = 10^−3^ for various coverages. The specific value of *θ* = 10^−3^ was chosen to reflect the median heterozygosity in a modern, non-African human individual.

We found the median of our *θ* estimates across replicates to be very close to the true value, but the variance to be a function of coverage. At low coverage (< 1x), *θ* was often overestimated, or inferred as zero. This is not surprising as the information about genetic diversity can only come from sites covered at least twice, which is rare at average coverages < 1x. As soon as average coverage exceeded 1.5x, however, our approach estimated *θ* at 10^−3^ very accurately (Fig. 1A).

**Figure 1.**
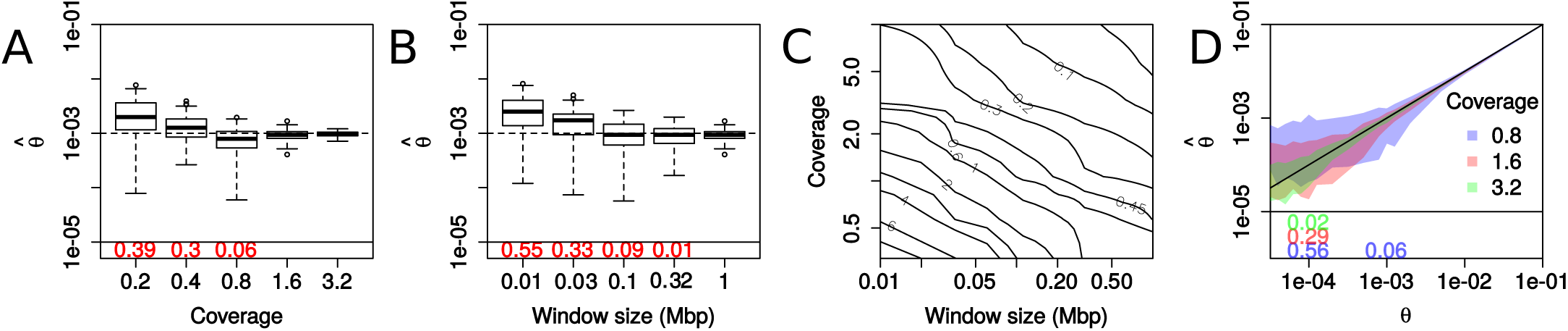
Power to infer *θ* from low coverage data. Results from sets of 100 simulations with post-mortem damage for different average coverage, window size and true *θ* values. (A) Estimated 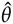 in windows of 1 Mb as a function of average coverage. (B) Estimated 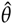 as a function of window size and fixed average coverage of lx. (C) Accuracy of estimating *θ* = 10^−3^ quantified as the median relative error 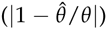 over replicates indicated by contour lines as a function of both coverage and window size. (D) True versus estimated *θ* for different average coverages (see color legend). Polygons indicate the 95% quantile of estimated 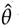 values among all replicates. The diagonal black line indicates the expectation for perfect estimation. In panels A,B and D, replicates resulting in a 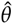 < 10^−5^ are not shown, but their percentage across replicates are printed below the horizontal black line.

We next performed simulations with a fixed coverage of 1x, but varying the window size (Fig 1). Interestingly, we found that an increase in window size has a positive effect on the estimate accuracy, similarly to an increase in coverage, suggesting that larger windows help to increase accuracy if coverage is very low. To illustrate this effect, we performed simulations at various window sizes and coverages and recorded the relative estimation error for a series of replicates. As expected, we found the median relative estimation error to be a direct function of the product of window size and coverage (Fig 1C), thus suggesting our method to perform well also at average coverages below 1x if the window size is large enough.

Using a third set of simulations, we found that at equal coverage and window size, higher *θ* values are estimated more accurately than lower values (Fig. 1D). This is expected since in the case of low *θ*, only few heterozygous sites are present in a given window, rendering the estimate more dependent on the detection of individual sites. Nonetheless, we found our approach to infer *θ* > 10^−4^ very accurately in a window of 1Mb if the average coverage exceeds 3x.

All results above were generated assuming base-specific quality scores to be normally distributed with *μ*_*Q*_ = 20 and standard deviation *σ*_*Q*_ = 4.5, which is the minimum quality expected with current sequencing approaches. Sequences generated with higher quality will positively affect estimation accuracy. Indeed we found that simulating data with *μ*_*Q*_ = 40 or *μ*_*Q*_ = 60 resulted in much lower estimation error, effectively rendering the estimation of *θ* feasible even at very low average coverage (2).

**Figure 2.**
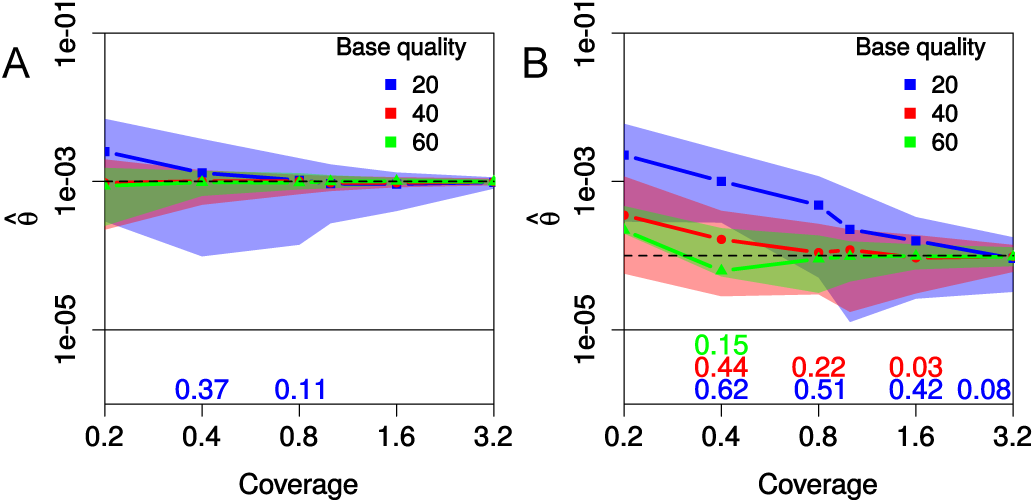
Effect of sequencing quality on power to estimate *θ*. Results from sets of 100 simulations to assess the power to estimate *θ* of 10^−4^ and 10^−3^ for panels A and B, respectively, for different average base qualities distributed normally with mean 20, 40 or 60 and a standard deviation of 4.5, but truncated at 0. Polygon shapes indicate the 95% confidence interval for estimated 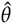 over all replicates, excluding those resulting in 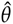 < 10^−5^ (the percentage excluded across are printed below the horizontal black line). All simulations were conducted with PMD and the true PMD probability functions were used during the estimation.

For instance, we found that at an average coverage of 0.8x, more than 90% of windows with *θ* = 10^−4^ and *μ*_*Q*_ = 60 were estimated within less than half an order of magnitude from the true value. At *μ*_*Q*_ = 20, this accuracy was only reached with an average coverage of 3.2x.

### Accuracy of Recalibration

The results discussed so far were all obtained under the assumption that quality scores provided by the sequencing machine are accurate. Unfortunately, this is rarely the case, making recalibration of the quality scores necessary for most applications, and in particular when trying to infer genetic diversity from low coverage data. Here we developed an approach to recalibrate quality scores without prior knowledge of the underlying sequencing information. Instead, we simply assume that a part of the sequence is known to be monomorphic, such as for instance the haploid X-chromosome in mammalian males.

To investigate the power of our approach to infer recalibration parameters, we simulated sequencing reads from a haploid region where the quality scores provided in the SAM files were distorted. We did this by first simulating fake quality scores from a uniform distribution *U*[5,60] and then transforming them into true quality scores according to eq. 12. We used the following coefficients: all context coefficients = 1.0, the coefficients for the raw quality score *β*_*q*_ = 1.5, the square of the raw quality score *β*_*q*^2^_ = 0.05, the position within the read *β*_*p*_ = −0.1 and the square of the position within the read *β*_*p*^2^_ = 5 · 10^−5^. These values were chosen to reflect a distortion observed in real data from the ancient human samples analyzed in this study (see below). They also result in both a relatively strong distortion as well as decent error rates for the evaluation of our approach.

We found all coefficients to be inferred with high accuracy from a 1Mb window with an average coverage above 1x (Fig. 3). If the amount of data was much lower than that, estimates were generally less accurate. In particular, we found the coefficients for the quality (*β*_*q*_ and *β*_*q*^2^_) to be often slightly overestimated at low coverages, likely because many sequencing errors go undetected since they can only be inferred at sites covered at least twice. However, this bias can be alleviated with larger window sizes if coverage is very low (see below).

**Figure 3.**
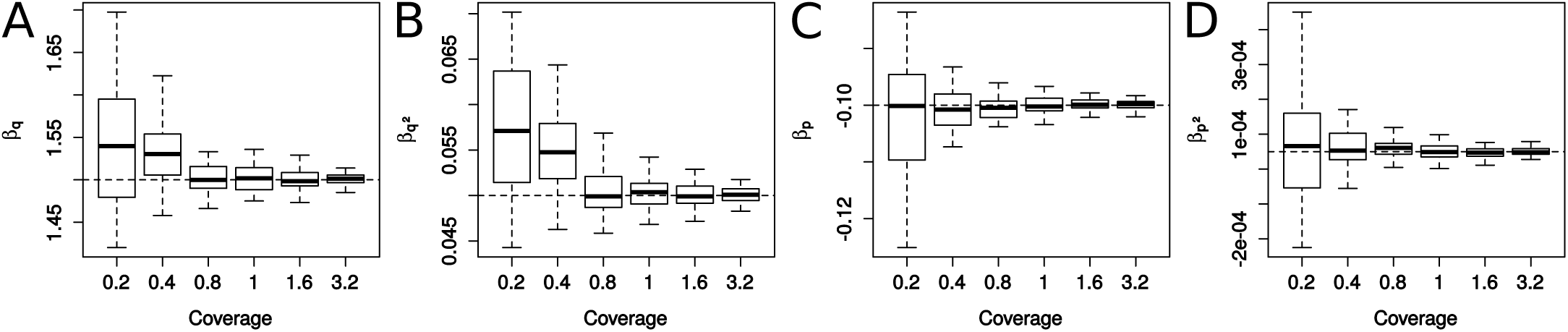
Accuracy in inferring recalibration parameters. Results from sets of 100 simulations are shown where sequence data from a haploid 1Mb region was simulated assuming a uniform distribution of observed quality scores (*U*[5,60]) that were then transformed to true qualities according to eq. 12 with *β*_*q*_ = 1.5,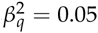,*β*_*p*_ = −0.1, 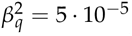 and all context coefficients at 1.0. All simulations were conducted with PMD and the true PMD probability functions were used during the estimation.

### Accuracy of full pipeline

We finally used simulations to assess the accuracy of the full pipeline, that is, when inferring first the pattern of PMD, then the recalibration coefficients given the inferred PMD pattern, and lastly using the recalibrated quality scores along with the inferred PMD pattern to estimate *θ*. In these simulations, the distortion of quality scores was, in addition to the four effects included above (*β*_*q*_ = 1.5, 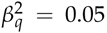, *β*_*p*_ =-0.1 and 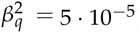), also affected by sequence context in that simulated sequencing errors were 1.5 times more likely to result in a *C* or *G* than in an *A* or *T*.

Regardless of the true *θ* value we used, we detected a strong bias in our estimates whenever very little data was used (Fig. 4). This is a direct result of the overestimation of the quality scores during the recalibration step as reported above, which leads to an overestimation of diversity. Encouragingly, however, this bias is overcome with only slightly more data. Indeed we found 1Mb of data with an average coverage of well below 1x to be sufficient to accurately infer *θ* ≥ 10^−3^ and pf 1x for *θ* = 10^−4^. Notably, even lower average coverages were sufficient when data was available for 10Mb. Finally,we found an average coverage of 4x to be sufficient when conducting recalibration and inference in windows as small as 0.1Mb. These results thus suggest that our approach may be useful not only for hemizygous individuals with large chunks of haploid DNA (the sex chromosomes), but may also work well in other individuals when using mtDNA or ultra conserved elements for recalibration.

**Figure 4.**
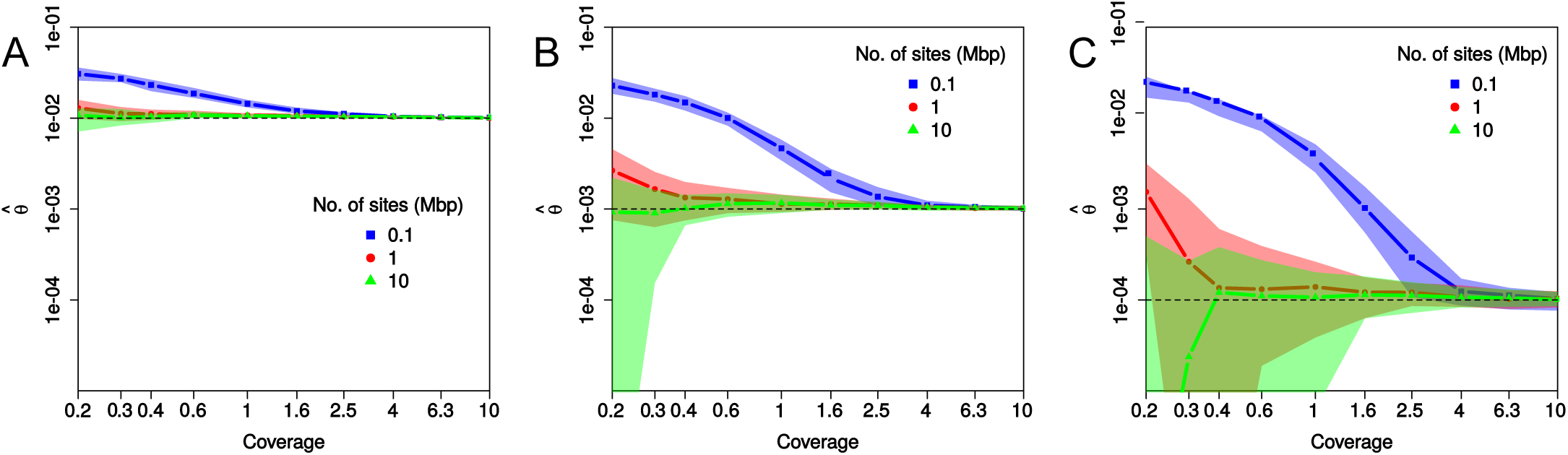
Accuracy in estimating *θ* using the full pipeline. Results from sets of 50 simulations each consisting of data from a haploid as well as a diploid region used to conduct recalibration and inference of *θ*, respectively. The data sets in panels A, B and C were simulated with different true values of *θ*, which are indicated with the dashed lines and were 10^−2^,10^−3^ and 10^−4^, respectively. Each data set was simulated with PMD as well as distorted base quality scores according to eq. 12 with *β*_*q*_= 1.5,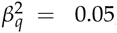,*β*_*p*_ = −0.1,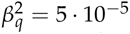. In addition, these simulations also included context effects in that sequencing errors were simulated to result 1.5 times more often in a *C* or *G* than in an *A* or *T*. The average coverage indicated is for the diploid data, while the hap-loid data was simulated with half the coverage. Line segments and polygons correspond to the median and the 90% quantile of all estimated 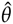 within the set of simulations, respectively.

## Application

We illustrate the benefit of our approach by inferring *θ* for several ancient human male samples and comparing these estimates to those obtained for several male individuals from the 1000 Genomes Project. For the ancient genomes we first inferred PMD patterns using the exponential model introduced here, then used the first 20Mb of the X chromosome to perform recalibration individually for each read group, taking the inferred PMD pattern into account. Finally, we used both the inferred PMD patterns as well as the recalibrated quality scores to infer *θ* in windows of 1Mb in the whole genome, excluding windows closer than 5Mb to Telomeres or Centromeres as defined by the track Gap in group *Mapping and Sequencing* in the UCSC Table Browser (Karolchik *et al.* 2008). The samples that we analyzed this way were 1) two European hunter-gatherer individuals (Jones *et al.* 2015), namely the Mesolithic genome “Kotias” from Kotias Klde cave from Western Georgia (KK1), and the western European Late Upper Palaeolithic genome, “Bichon” from Grotte du Bichon, Switzerland (Bich), approximately 17.700 years old 2) an individual from the Bronze age burial site at Ludas-Varjú-dülö, Hungary (BR2 Gamba *et al.* 2014) and 3) a 4500 years old male from Mota Cave in the southern Ethiopian highlands (Mota Gallego Llorente and Jones 2015). All these samples had relatively high coverage (>10x) and thus allowed us to infer fine scaled patterns of heterozygous along the genomes, even for regions with low diversity (*θ* < 10^−4^).

For comparison, we also inferred diversity patterns for nine modern males from three populations that were analyzed as part of the 1000 genomes project phase 3 (alignment files downlaoded from ftp://ftp.1000genomes.ebi.ac.uk/vol1/ftp/phase3/data/). These were the British males HG00115, HG00116 and HG00117, the Tuscany males NA20509, NA20511 and NA20762, and the Yoruban males NA18486, NA18519 and NA18522. As shown in Fig. 5A, these nine individuals portray the expected pattern of higher diversity in African than European individuals, but they also revealed significant variation among individuals of the same population (t-test, *p* < 10^−5^ in at least 2 out of 3 possible comparisons in each population). Larger differences in overall diversity was observed among the ancient samples analyzed. Unsurprisingly, the African sample Mota exhibited the highest diversity of all ancient samples, which was also higher than the diversity observed in modern day Europeans, yet lower than modern day Yorubans. The ancient sample with the second highest diversity was the Bronze Age sample BR2, whose diversity falls well within the range of estimates obtained from modern day Europeans. In contrast, the two European hunter-gatherer samples KK1 and Bichon showed much lower diversity than modern day Europeans with their median estimates being 15-25% lower than the median estimates of modern Europeans. These results thus suggest that while hunter-gatherer populations had much lower diversity, the diversity found in Europeans about 3,000 years ago was very comparable to the diversity observed today. This conclusion is in perfect agreement with with a temporal trend in the total length of runs of homozygoisty (ROH) inferred from imputed genotypes among ancient samples from Hungary that spanned a period from 5,700 - 1,000 BC and also included the sample BR2 (Gamba *et al.* 2014).

**Figure 5.**
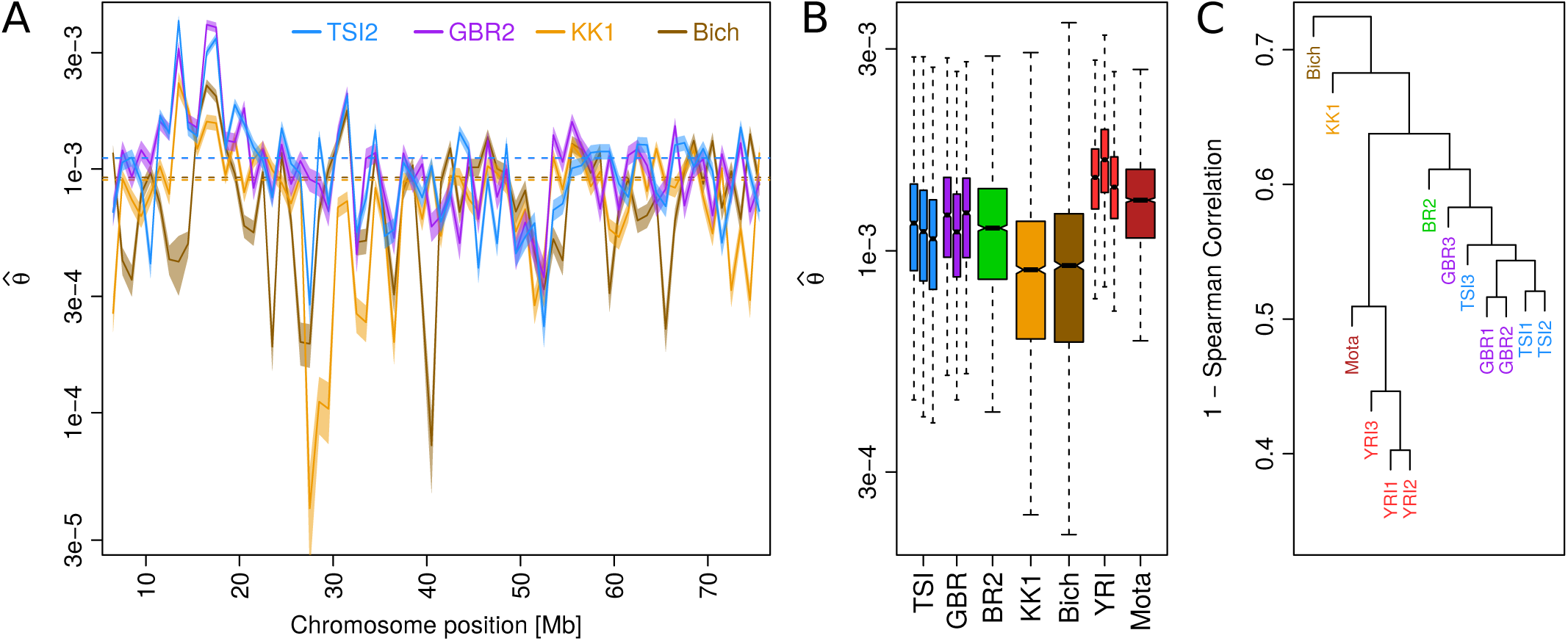
Local diversity in ancient and modern humans. A) Heterozygosity (*θ*) inferred in 1Mb windows along the first 75 Mb of chromosome 1 (excluding windows closer than 5Mb of the telomere) for two modern Europeans (TSI2 and GBR2) and two ancient European hunter-gatherers (KK1 and Bich). Solid lines indicate the MLE estimate, shades indicated the 95% confidence intervals and dashed lines the genome-wide median for each sample. B) Distribution of estimates 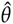 in 1Mb windows across the first 22 chromosomes of each sample. C) Similarity in the pattern of *θ* along the genome visualized by hierarchical clustering using 1-Spearman correlation as distance.

The inference of local diversity patterns also allows us to compare the distribution of diversity in the genome between individuals, regardless of the overall level of diversity. This analysis revealed a substantial phylogenetic signal in the distribution of diversity as quantified by Spearman correlations. For instance, the diversity pattern is more strongly correlated among modern Yorubans (Spearman correlations between 0.55 - 0.60) than between Yorubans and Europeans (0.40 - 0.50). Similarly, the diversity pattern of the ancient African samples Mota is most strongly correlated with that of modern day Yorubans (0.491 - 0.537), and much less so with modern day Europeans (0.362 - 0.433). Interestingly, European samples are more diverse in their patterns than Africans (Spearman correlations between 0.42 - 0.48) and their pair-wise correlations do not exceed those obtained when comparing African and European individuals. Nonetheless, hierarchical clustering groups all modern day Europeans together and also puts the Bronze age sample BR2 at the basis of that clade (Fig. 5B).

The lowest pair-wise correlations (0.28 - 0.39) were found in comparisons involving the two European hunter-gatherer samples KK1 and Bich, with the overall lowest being the correlation between these samples (0.28). This is also illustrated visually when plotting our estimates of the first 75Mb of chromosome 1 where we found relatively high concordance in local diversity among the two European samples, but vastly different patterns among the hunter-gatherer samples (Fig. 5C). These results suggest that despite very comparable overall levels of diversity, the distribution of diversity along the genome was very diverse among European hunter-gatherer populations and very different from the one observed among modern day individuals. Multiple observations support such a conclusion: first, the two samples analyzed here represent two vastly different geographic regions, with one being samples in Switzerland and the other in Georgia, and were previously reported to belong to two different clades that split 45,000 years ago as inferred from genotyping data (Jones *et al.* 2015). Second, the ancestry of modern Europeans traces only partly back to European hunter-gatherers with early Neolithic people from the Aegean (Hofmanová *et al.* 2015) and Yamnaya steppe herders (Haak *et al.* 2015) contributing the majority of the modern day genetic make up. Finally, the two European hunter-gatherer samples both exhibit many but unique regions of very low diversity (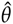 < 10^−4^ in 4% of all windows, compared to 0.00 - 0.03% in all modern Europeans), likely the result of small population sizes with some level of consanguinity in the population (Pemberton *et al.* 2012).

## Discussion

Quantifying genetic diversity and comparing it between different individuals and populations is fundamental to understanding the evolutionary processes shaping genetic variation. Unfortunately, the inference of heterozygosity is confounded by both sequencing errors resulting in false diversity as well as the statistical power to identify heterozygous sites, particularly when coverage is low. Several methods have been developed to learn about heterozygosity probabilistically, that is, without the need to first call genotypes. A rather recent such approach (Bryc *et al.* 2013) proposes to leverage data from external reference individuals to obtain an unbiased estimate of the probability that a specific sites is heterozygous. The expected heterozygosity is then estimated from these site specific estimates. Since this approach requires per site estimates to be accurate, only sites with a coverage of 5x or higher can be included in the analysis, which severely limits the scope of the application.

An alternative is to infer heterozygosity probabilistically from a collection of sites. Among the earliest such methods was a likelihood based estimator (Lynch 2008), which infers heterozygosity of an individual jointly with the rate sequencing errors. We presented a natural extension of this approach that relaxes the infinite sites assumption and integrates post-mortem damage (PMD), a particular feature of ancient DNA. We then showed that this allows for unbiased estimates a much lower coverage than the original estimator, which was found to be ill-behaved at coverages below 4x (Lynch 2008).

A downside of our approach is that it assumes sequencing errors to be known, while the original approach estimates the rate of sequencing errors jointly with heterozygosity. This, however, allows us to relax the assumption of constant error across all reads and to benefit from the quality information provided by current sequencing technologies. Yet since these provided quality scores are often distorted, we also introduced here a method to recalibrate the quality scores for low coverage genomes. In contrast to commonly used methods for recalibration (e.g. DePristo *et al.* 2011; McKenna *et al.* 2010; Cabanski *et al.* 2012), our approach does not require information about the underlying sequence context. It only assumes sites to be monomorphic while integrating over the uncertainty of the sequence itself. Examples of regions known to be monomorphic are the sex chromosomes in hemizygous individuals. But since we found that our method recalibrates quality scores with high accuracy and reliably even based on DNA stretches as short as 1Mb, we are confident that it will work even on ultra conserved elements or plasmid DNA. Finally, we note that if multiple individuals are sequenced together, they are likely affected by the same distortion of quality scores and can hence be recalibrated with parameters inferred from a subset of them (e.g. the male samples).

As an illustration, we applied the here developed methods to modern and ancient human samples of various coverage. While our approach to infer heterozygosity incorporates the possibility of PMD, it assumes that the probability of a PMD event occurring is known. We thus also introduce two methods to infer these probability functions from raw data that are robust to divergence between the sample and the reference genome. By inferring PMD patterns for each sample, then the recalibration parameters, and finally local diversity in 1Mb windows, we found that both ancient and modern African samples exhibited much larger diversity than European individuals. In addition, the diversity inferred from two ancient European hunter-gatherer samples was much lower than that of modern samples, which is likely explained by smaller population sizes. Besides overall differences in diversity, also the pattern of diversity along the genome revealed a strong geographic clustering among modern and ancient samples. The exceptions were the two European hunter-gatherers that showed patterns very different from both modern samples as well as from one another, further corroborating the view (Jones *et al.* 2015) that these samples represent different and ancient clades that contributed only marginally to the genetic make-up of modern day Europeans.

## Appendix Emission probabilities in the presence of post-mortem damage

Following Lynch (2008) and commonly used approaches (Li 2011, e.g.), we assume here that a sequencing read is equally likely to cover any of the two alleles of an individual and that sequencing errors may result in any of the alternative bases with equal probability *ϵ*_*ij*_ /3. In the absence of post-mortem damage (PMD), the probability of observing a base *d*_*ij*_ given the underlying genotype *g*_*i*_ = *kl* is then given by

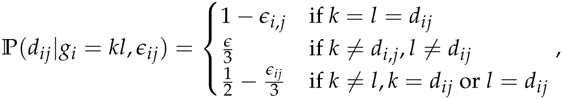

In ancient DNA, differences between the base observed within a read and the underlying alleles may also be the result of PMD. Let us denote by *D*_*C*→*T*_(*q*_*ij*_) and *D*_*G*→*A*_(*q*_*ij*_) the known probability that a *C* → *T* or *G* → *A* PMD occurred at the position covering site *i* in read *j*, respectively. In the presence of PMD, the probability of observing a base *d*_*ij*_ given the underlying genotype *g*_*i*_ = *kl* is given by

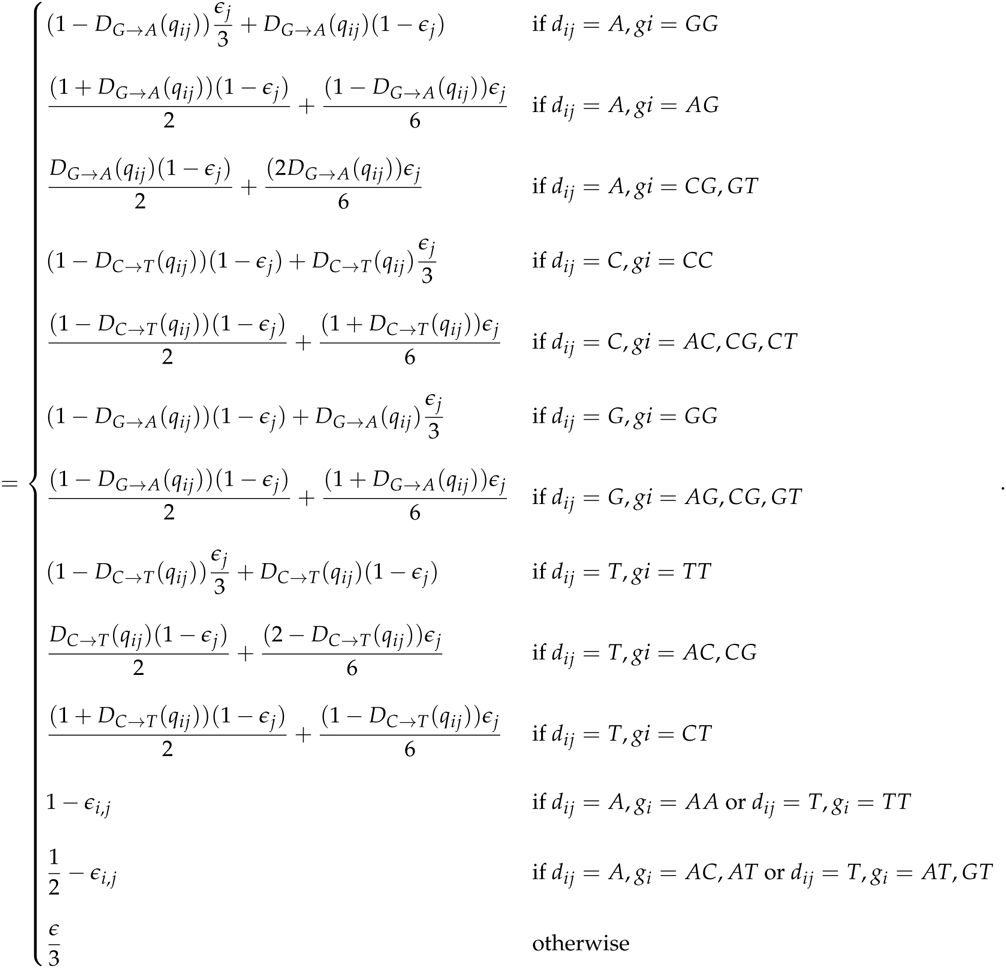

***Implementation of the EM algorithm to infer*** *θ*

Here we present an efficient implementation of the algorithm to infer *θ* for a genomic region containing I sites:

1. Calculate the matrix of emission probabilities 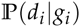 for all positions and genotypes according to the formulas given in the Appendix.
2. Estimate the initial base frequencies *π* from the base frequencies among all reads in the window.
3. Set the initial *θ* to the genome-wide expectation.
4. Run the EM algorithm by repeating the following steps until convergence is reached:

a. Set *θ*′ = *θ*, *π*′ = *π*, *ρ*′ = *e*^-*θ*′^ / (1 - *e*^-*θ*′^) and *μ*′ = 0.
b. Calculate substitution probabilities 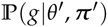 for all genotypes *g* according to eq. 1 using the current estimates of *θ*′ and *π*′.
c. Calculate 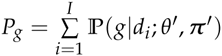 for all genotypes *g* according to eq. 3.
d. Find the new estimates of the parameters *θ* and *π* using the Newton-Ralphson method by setting **x** = {*π*′,*ρ*′, *μ*′} and repeating the following steps until convergence:

i. Calculate vector *F* according to eq. 4.
ii. Calculate **J**^−1^ according to eq. 6.
iii. Update **x** according to eq. 7.
e. (e) estimate new parameter estimates as *π*_*k*_ = *x*_*k*_ and *θ* = −log(*x*_5_ /(1 + *x*_5_)).

